# The SPARC DRC: Building a resource for the autonomic nervous system community

**DOI:** 10.1101/2021.04.01.438136

**Authors:** Mahyar Osanlouy, Anita Bandrowski, Bernard de Bono, David Brooks, Antonio M. Cassarà, Richard Christie, Nazanin Ebrahimi, Tom Gillespie, Jeffrey S. Grethe, Leonardo A. Guercio, Maci Heal, Mabelle Lin, Niels Kuster, Maryann E. Martone, Esra Neufeld, David P. Nickerson, Elias G. Soltani, Susan Tappan, Joost B. Wagenaar, Katie Zhuang, Peter J. Hunter

## Abstract

The Data and Resource Center (DRC) of the NIH-funded SPARC program is developing databases, connectivity maps and simulation tools for the mammalian autonomic nervous system. The experimental data and mathematical models supplied to the DRC by the SPARC consortium are curated, annotated and semantically linked via a single knowledgebase. A data portal has been developed that allows discovery of data and models both via semantic search and via an interface that includes Google Map-like 2D flatmaps for displaying connectivity, and 3D anatomical organ scaffolds that provide a common coordinate framework for cross-species comparisons. We discuss examples that illustrate the data pipeline, which includes data upload, curation, segmentation (for image data), registration against the flatmaps and scaffolds, and finally display via the web portal, including the link to freely available online computational facilities that will enable neuromodulation hypotheses to be investigated by the autonomic neuroscience community and device manufacturers.

## 1 INTRODUCTION

The physiological function of every visceral organ in the body is linked intimately with autonomic control, and dysfunction of the autonomic nervous system (ANS) is often associated with chronic disease in these organs (Low, 2011). The NIH Common Fund’s ‘Stimulating Peripheral Activity to Relieve Conditions’ (SPARC) program is aimed at providing an anatomically and physiologically accurate description of the ANS in order to assist in the design and testing of medical devices that can modulate the ANS for therapeutic benefit^1^. A primary motivation for the project was the realization that designing therapeutically effective neuro-modulation strategies for bioelectronic medicine is extremely difficult without a quantitative understanding of the very complex connectivity and function of the ANS.

The SPARC program is organized as a consortium, with investigators funded to obtain data on organ-ANS interactions and the SPARC Data and Resource Center (DRC) to provide the infrastructure for data and tools. The DRC has four separately funded components or DRC ‘cores’, including academic, not-for-profit, and commercial organisations. The DRC cores store and process the data from the SPARC experimental community: (i) DAT-Core is providing data storage, management, and publication platforms; (ii) K-Core is providing a workflow for data curation and annotation, and a knowledgebase of neural connectivity and function; (iii) MAP-Core is providing tools for data processing, including image segmentation, and is mapping disparate data sets from multiple species into a common coordinate framework; and (iv) SIM-Core is providing a platform for computational modeling of the role of the ANS in regulating physiological activity and of neural interfaces and their neuromodulatory impact, based on the mapped pathways and computational models of ANS and organ function developed by SPARC modelers. Together these cores work to create an integrated environment for exploring data and knowledge on the ANS; coupling them to a web portal (https://sparc.science) for accessing these data and for access to a sophisticated simulation platform ^2^. Figure 1 illustrates the workflow for data upload, curation, registration, and publication.

**Figure 1.**
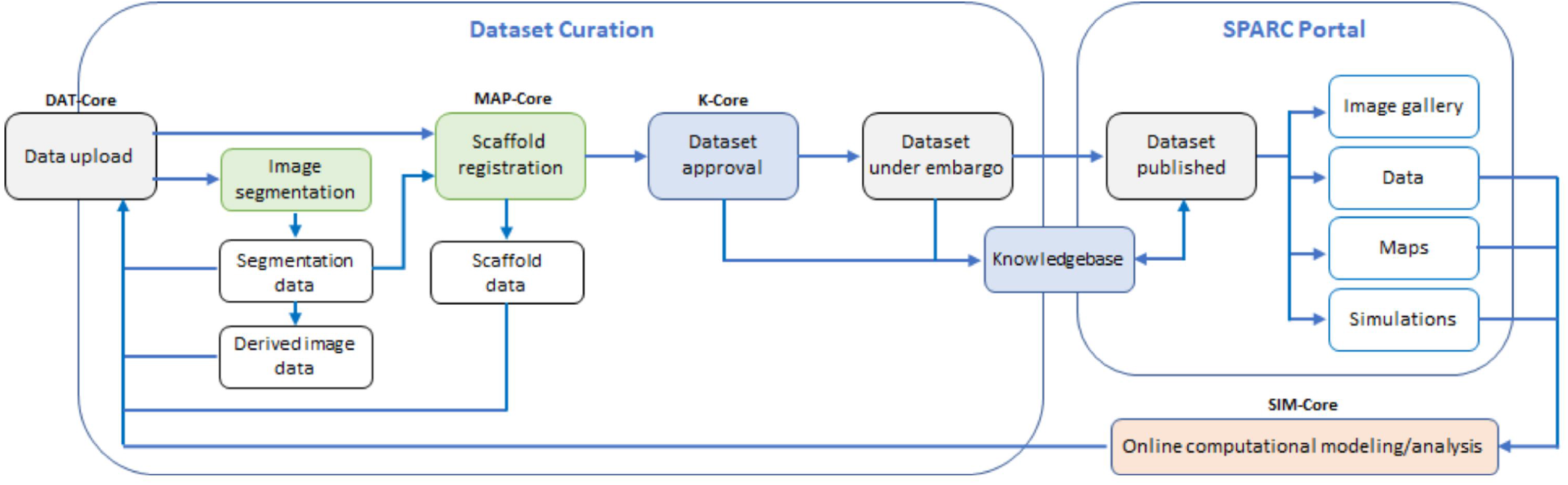
The data workflow: Experimental data and mathematical models created by the SPARC community are uploaded to a data repository managed by DAT-Core, along with metadata that is verified by K-Core and used to create a semantic knowledgebase for the ANS community. Image segmentation and other data processing steps, including mapping data and models into appropriate anatomical locations using common coordinate systems, is undertaken by MAP-Core. The simulation environment for using the data and models to understand ANS function, and to assist in the development of neuromodulation therapies and devices that modulate that function, is provided by SIM-Core. All cores contribute to the https://sparc.science portal that provides a resource for the ANS biomedical science and bioengineering communities, and for the manufacturers of neuromodulation devices.

In this paper we review the contribution that the DRC is making to the neuroscience goals of the SPARC program. Many of the methods and open source infrastructure being developed for SPARC may also be of interest to the broader audience of biomedical scientists interested in the reproducibility of experimental data and the ability to compare data across species, and who also have an interest in mapping their data into quantitative standards-based modeling frameworks.

The types of experimental data being generated by SPARC investigators are primarily: (i) the connectivity between visceral organs and the ANS for a given species; (ii) neural pathway image data from a wide variety of modalities; (iii) electrophysiological data; (iv) motility data (deformation of the beating heart, inflation of the lungs, filling of the bladder, contraction waves in the gut, etc); and (v) RNAseq data from a variety of cell types. Computational models developed by SPARC can be mechanistic or data-driven and range from neural interface models, simulating the impact of physical exposure of neural and anatomical structures on electrophysiological activity, to organ physiology models, neural control models, data analyses, and machine learning.

In the following, we provide an overview of the SPARC infrastructure, starting with the SPARC Portal (Section 2). We then provide more detail about the data management platform (Section 3), the curation and annotation of SPARC data (Section 4), the processing of neural connectivity data (Section 5), image segmentation and annotation (Section 6), and common coordinate systems and 3D scaffolds for data mapping (Section 7). Examples showing how organ data are processed through this pipeline are given in Section 8. The incorporation of data and models into the computational framework is described in Section 9. In Section 10 we discuss our plans for further developments of the portal.

## 2 THE SPARC PORTAL

The SPARC Portal (https://sparc.science), which is being developed jointly by the four cores of the SPARC DRC, is the central access point for information on and from the SPARC initiative. It provides a unified interface for users to interact with SPARC resources and when fully developed, will provide a coherent user workflow around public data generated by the SPARC investigators. Note that FAIR principles are adopted, to ensure that data is ‘findable, accessible, interoperable and reusable’^3^ (Wilkinson et al., 2016), and all software developed by the project is released under an open source license.

The core functions of the SPARC Portal are to:

1. provide information about the SPARC program, outreach, and events
2. allow users to discover, find, access and visualize curated public datasets generated by the SPARC consortium
3. allow users to see these data in the context of their anatomical location and to visualize and search anatomical and functional relationships
4. leverage these data to launch analyses and simulations on o^2^S^2^PARC that provide insight towards understanding and influencing the autonomic nervous system and its role in regulating organ physiology.

The SPARC Portal is a front-end web-application that provides a view into several underlying platforms, such as *Blackfynn Discover*^4^, *UCSD’s SciCrunch Platform*^5^, *MBF Bioscience Biolucida*^6^, and *IT’IS o*^2^*S*^2^*PARC*^7^, that deliver content for the application. These are all discussed further below.

The vision for the targeted functionality of the portal is to:

1. enable visitors to explore and discover SPARC output (data, resources, publications, documentation, events, etc.) and in some cases making use of dedicated viewers to inspect data without downloading it (e.g. the *Biolucida* for microscopy images)
2. provide up-to-date information and insight about/into the ANS and its integrated physiological role (e.g., through consolidated and curated maps)
3. facilitate collaboration between SPARC researchers on SPARC-related topics
4. facilitate finding of related/associated data, computational models, analysis functionality, and anatomical models
5. support FAIRness in data/model publication (based on the curation work and the FAIRness components of other DRC resources)
6. serve as a gateway for SPARC communication (internal and external; incl. news and events) and community building.

Experimental data acquisition and computational modeling are tightly connected, in SPARC and science in general. Experimental data serves to inform, feed, and validate computational models, while computational models help formulate testable hypotheses, interpret experimental data, and provide dense information under highly controlled conditions. Computational modeling also plays a central role in producing derived data from primary measurement data. Hence, similar concerns about integration, sustainability, and FAIRness apply to computational modeling. Achieving integration, sustainability, and FAIRness of computational modeling and data analysis comes with its own set of specific challenges:

1. computational models are developed using a wide range of techniques and platforms (e.g., Matlab and Python scripting, C++ programing; HPC-enabled physics simulators, machine learning-based models, analytical models) – this variability a priori prevents integration and coupling of these models as part of a larger model
2. the implementation of computational models is typically platform-specific – it is hard to ascertain that it reliably produces (the same) results on another machine, operating system, compiler, library version, thus encumbering reproducibility, maintenance and sharing
3. the process by which derived data is produced from original (e.g., measurement) data is rarely well documented and preserved – e.g., it is not published along with scientific papers – hampering reproducibility
4. similar analysis routine therefore gets reimplemented repeatedly, or previously developed analysis scripts are adapted without keeping track of provenance, changes, and variants
5. academically developed software is frequently poorly documented, has not been developed with the degree of testing and quality assurance required, e.g., for regulatory purposes, and does not possess user-friendly (graphical) user-interfaces
6. these points frequently result in a lack of long-term usability of scientifically developed software and models and associated loss (e.g., once the corresponding PhD has left the lab)
7. similarly, establishing workflows that combine usage of multiple different tools represents a form of expertise that is hard to preserve
8. it is difficult to gauge the degree to which computational models have been validated and thus, the extent of their reliability and their applicability range
9. while academic research is frequently published, as publications are a common success metric and proper credit is attributed through citations, the open sharing of computational tools is not similarly established and encouraged.

How all of these challenges are addressed in SPARC, e.g., through an online accessible and integrative computational platform, is discussed in Section 9. Online hosting of computational models and analysis tools also results in benefits, such as facilitation of joint development, the possibility of providing a demonstrator (e.g., as supplementary material to a scientific publication, or a grant application), and access to computational resources (e.g., for high-performance-computing or large memory requirements). In the following sections, we go into more detail about the components of the SPARC DRC platform.

## 3 DATA MANAGEMENT AND DISTRIBUTION

A scalable cloud-based platform for data storage, management and distribution has been developed in partnership with the SPARC program and is leveraged by the SPARC investigators to upload, share, and publish datasets associated with the SPARC program. The Blackfynn Data Management (DMP) and Discover platforms^8^ provide the technical infrastructure to support the curation and data publication processes within the SPARC program. The platform was developed to provide a sustainable [10] solution for managing and securely distributing terabyte-scale datasets.

The DMP was developed from the beginning with the notion that public scientific datasets need to adhere to FAIR principles of data sharing^9^ and provide mechanisms to make very large datasets readily available for downstream analysis in the cloud. Rather than focusing on providing a data repository for files, the DMP provides advanced support for complex metadata describing the datasets. By integrating support for files and metadata, the platform enables scientists to publish high quality datasets that can be used to generate derived work. This will ensure that the SPARC Data Core not only ameliorates the “hard drive” problem of scientific research [11], but also facilitates easy data sharing and collaborative research.

The DMP and Discover platforms are FAIR compliant and implement all recommended functionality to ensure datasets are findable, accessible, interoperable, and reproducible (Wilkinson et al., 2016). This includes associating globally defined identifiers (DOIs) with each version of a dataset on the Discover platform, integration with ORCID, ensuring that dataset metadata is represented on the platform for indexing purposes by Google Dataset and other web-crawling platform, and the adoption of FAIR vocabularies and community standards. Given the size of SPARC datasets (sometimes in the TB range), special consideration is given to enable users to analyze and access the data in the cloud in a sustainable way as we envision the direction of scientific progress to increasingly rely on data analysis in the cloud, over a model where data is downloaded and processed locally.

The process of publishing a dataset is a multi-step process. First, the data are uploaded and curated with the help of the SPARC Curation team. When a dataset is ready to be distributed, the SPARC PI requests that the data is published (and optimally specifies a release date). A publication team provides a final review of the dataset and approves the publication request after which a snapshot of the dataset is created and published to the Discover platform. Once published, anybody can find/access the public dataset. If the dataset is published with a release date, anybody can find the dataset, but early access needs to be requested from the dataset owner.

The DMP provides robust programmatic access through its open API, command line interface, and Python client. These API’s are leveraged by a number of other SPARC initiatives such as the SPARC Portal, Biolucida, o^2^S^2^PARC and the SPARC SODA Tool^10^ (see below).

## 4 DATA CURATION PROCESSES, ANNOTATION AND KNOWLEDGE MANAGEMENT

### 4.1 Standardization Initiatives: data, models and simulations

Curation and knowledge management are provided through K-Core, led by the FAIR Data Informatics Laboratory at UCSD (FDIlabs.org). SPARC has designed its public data platform and curation standards to make all SPARC data FAIR. All data are accompanied by rich metadata, including descriptive metadata about the project and authors, a full experimental protocol and structured experimental metadata. Each published dataset receives a persistent identifier in the form of a Digital Object Identifier (DOI), and also includes full dataset citation information so that datasets can be formally cited (Fenner et al., 2019; Starr et al., 2015).

Each dataset produced by a SPARC investigator is subjected to automated and human curation. The curation process is shown in Figure 2. To date, SPARC has been curating data to two primary standards developed by the SPARC consortium: 1) The Minimal Information Standard (MIS), a semantic metadata scheme capturing key experimental and dataset details^11^; 2) The SPARC Dataset Structure (SDS), a file and metadata organizational scheme based on the Brain Imaging Data Structure (BIDS) developed by the neuroimaging community (Gorgolewski et al., 2016). SDS defines a common folder organization and file naming convention along with required and recommended metadata. Details about SDS can be found in (Bandrowski et al., 2021). SPARC investigators are required to submit their data in SDS along with a detailed experimental protocol deposited in Protocols.io^12^; SPARC curators then align the submitted metadata to the MIS using automated and semi-automated workflows. To assist researchers in organizing and annotating their data to SPARC, a submission tool has been developed, SODA (Software for Organizing Data Automatically)^13^ (Bandrowski et al., 2021).

**Figure 2.**
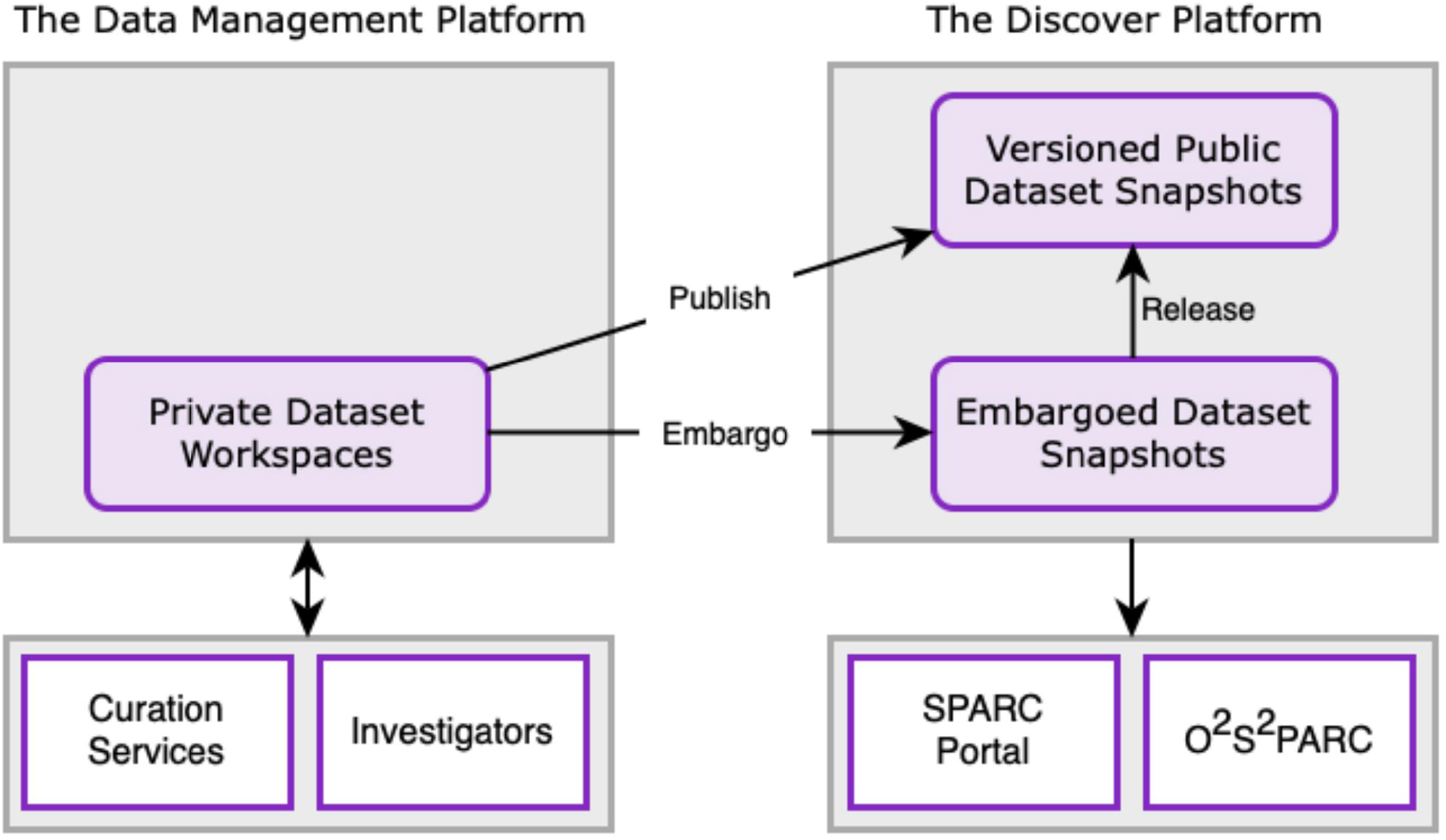
Schematic overview of the relationship between the DMP, Discover, SPARC portal and o^2^S^2^PARC applications. Data are uploaded to the DMP and curated through the SPARC curation workflow. Once a dataset is curated, a snapshot of the dataset is persisted in the Discover platform and a DOI is assigned. Initially, the data is released as an embargoed dataset, with a final release date on which the entire dataset is made publicly available. Users have the ability to request early access to embargoed data through the platform after signing a SPARC Data Use Agreement.

As the SPARC program progresses, additional standards will be implemented to increase the depth of FAIRness for our data and to bring SPARC into alignment with other large initiatives such as the US BRAIN Initiative^14^. Towards that end, additional standards for file formats and modality-specific metadata, e.g., imaging and physiological metadata, are in the process of being evaluated. SPARC has established a Data Standards Committee comprising researchers who review and approve all recommended standards. A microscopy imaging metadata standard was recently approved and an openly available application, MicroFile+^15^, has been created in order to align image data from many microscope sources with this standard.

SPARC data are aligned to common spatial standards through the processes described in Section 7, and also annotated with FAIR vocabularies derived from community ontologies like UBERON (Mungall et al., 2012). A list of ontologies currently employed in SPARC can be found in the SDS White Paper (Bandrowski et al., 2021). SPARC is enhancing these community ontologies with additional content relevant to SPARC, using the vocabulary services developed by the Neuroscience Information Framework^16^. These services include Interlex, an on-line database for storing, serving and building vocabularies^17^, and SciGraph, a neo4j-backed ontology store^18^ maintained by FDILabs^19^.

K-Core is also building and managing the SPARC Knowledge Graph (SKG), a semantic store that augments data sets described according to the MIS supplemented with knowledge about anatomical relationships. These relationships include partonomies encoded in ontologies and the SPARC Connectivity Knowledgebase, a comprehensive store of semantic knowledge about ANS connectivity. The SPARC Connectivity Knowledgebase is populated through the development of detailed circuitry based on SPARC data and expert knowledge using the ApiNATOMY platform (see Section 5) supplemented with knowledge derived from the literature. The SKG is managed using the SciGraph infrastructure. The SKG can answer queries such as “Find all neurons with axons that run through the pudendal nerve.” It powers the MAP interface in the portal and also the dataset descriptions available through the SPARC Portal (Section 2). In upcoming releases of SPARC, the SKG will be available for direct query.

### 4.2 Curation for computational modeling

In addition to the general curation issues (such as provenance, ownership, protocols documentation, dependencies, and versions) discussed above, curation of computational services, computational studies, and computationally derived results also have specific requirements, which are reflected in the recently introduced new computational MIS (cMIS). Computational services must have associated meta-data that documents the required inputs (type, format, etc.) and produced outputs (potentially even their physiological meaning), the hardware requirements, the degree of reliability (e.g., degree of verification, validation, and certification, suitable context-of-use - the quality assurance information is organized according to the ‘Ten Simple Rules’ (Erdemir et al., 2020)), etc. An online form is available for submitting the required information as part of the semi-automatic service-creation-process. Curation of derived data (results) obtained through computational modeling or analysis is simplified, first of all, because it originates from a computational study and input data that can be linked and referenced to ascertain reproducibility, and secondly, because the generation through a computational pipeline allows to automatize the attachment of a large part of the required meta-data (e.g., services know the units, dimensions and types of the results they produce, as well as the execution time and owner – a service might even know something about the physiological meaning of an output it produces). Similarly, computational studies are generated on o^2^S^2^PARC and consist of inputs, scripts, and services which are routed through the SPARC data repositories and hence traceable (full chain-of-custody) and reproducible. FAIRness for models and simulations is also achieved through active support of or compatibility with selected community standards. Particular consideration is paid by standards and initiatives from the Physiome project^20^ (Hunter and Borg, 2003), e.g., by storing references to computational models and scaffolds in the Physiome Model Repository^21^ and by offering dedicated services tailored to CellML models^22^. Note that CellML is an XML-based language to store and exchange computer-based mathematical models (Lloyd et al., 2004); o^2^S^2^PARC includes an OpenCOR solver (Garny and Hunter, 2015) for CellML models, as well as a service that facilitates the creation of new computational services based on CellML specifications.

## 5 A CONNECTIVITY MAP AND KNOWLEDGEBASE FOR THE ANS

A major challenge facing the SPARC effort is to enable researchers to navigate, cluster, and search experimental data about the ANS and viscera in terms of physiological relevance. We address some of these challenges through the use of the ApiNATOMY toolkit (De Bono et al., 2012) that supports data integration over a multiscale connectivity model of cells, tissues and organs.

As a framework, ApiNATOMY consists of a knowledge representation, and knowledge management tools, that enables topological and semantic modeling of process routes and associated anatomical compartments in multiscale physiology (see (De Bono et al., 2018) and Figure 3). Field experts (e.g. SPARC experimentalists) provide model specifications in a concise semi-structured format based on predefined templates designed to define common topological structures (such as neural arborizations, etc.). ApiNATOMY tools then extend these specifications to a fully operational graph that can be visually manipulated, serialized and integrated with other knowledge bases such as NIF-Ontology^23^ (Bug et al., 2008).

**Figure 3.**
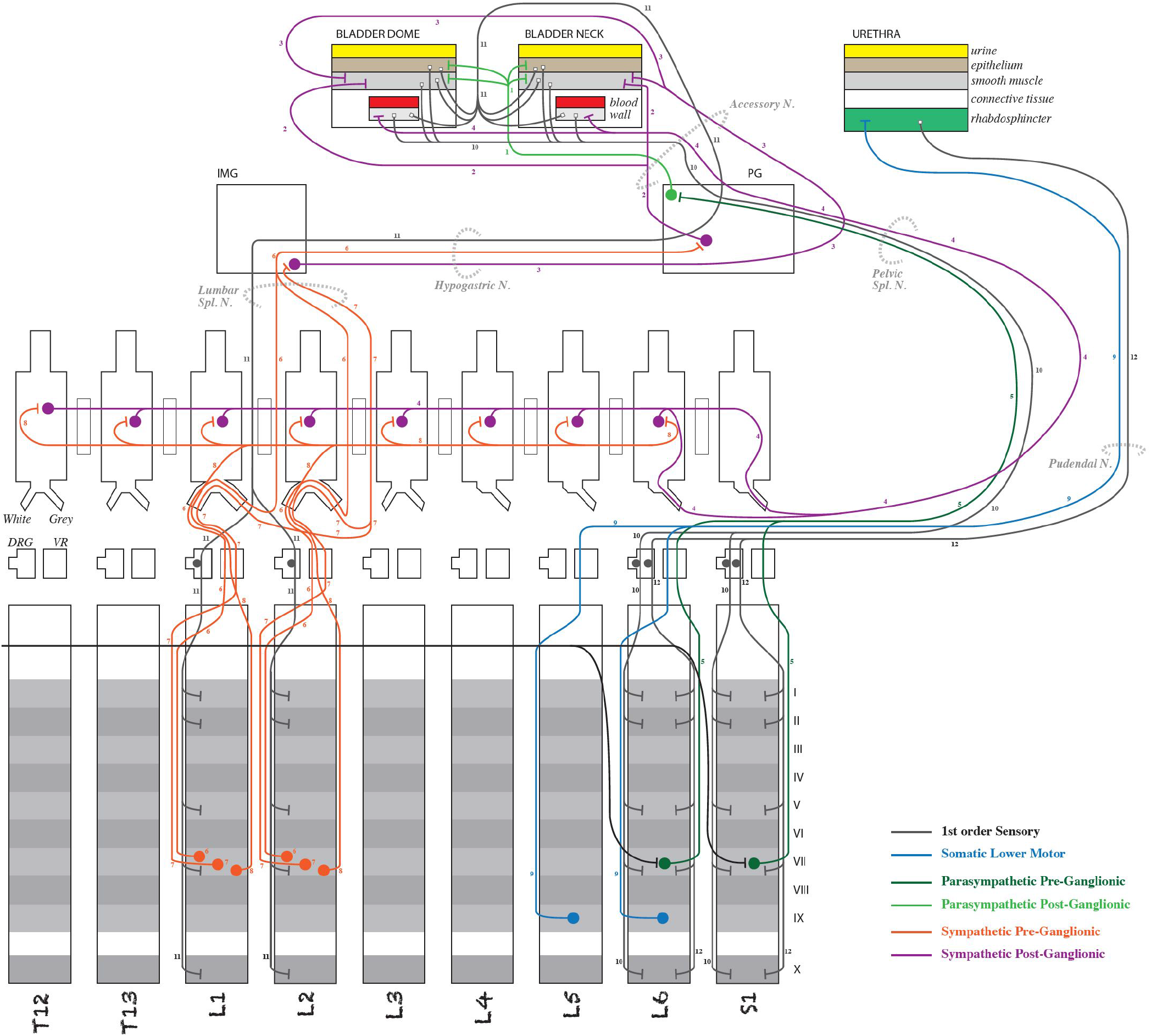
Detail of an ApiNATOMY schematic diagram representing neural connectivity between the rodent spinal cord and the lower urinary tract [Model available [here](https://github.com/open-physiology/apinatomy-models/tree/master/keast-bladder)]. Note that the pathway information in diagrams such as this is stored in SPARC Connectivity Knowledge Base and used to render the pathways in the more anatomically oriented flatmaps (see Figure 4)

As ApiNATOMY models are annotated to the same SPARC standard as MIS for experimental data, this connectivity knowledge is then leveraged to enhance navigation, classification and search for the Portal. For instance, if a SPARC researcher is designing a procedure on the Middle Cervical Ganglion (MCG), the Portal now can provide the means to discover existing SPARC data collected along the route of neurons passing through the MCG that might help with experimental planning.

The ApiNATOMY infrastructure provides an interactive graphical web application (http://open-physiology-viewer.surge.sh/) as well as the means to query linked metadata to discover datasets, track provenance, etc. To that end, the infrastructure supports a generalizable approach for converting application specific JSON data structures into RDF using JSON-LD. This approach has proved to vastly simplify the complexity of the system and has made it easier to maintain. These exports are incorporated into the SPARC Connectivity Knowledge Base.

The connectivity illustrated by the ApiNATOMY diagrams and transferred with the semantic annotation to the SPARC Connectivity Knowledge Base is then used algorithmically to generate flatmap diagrams of ANS connectivity for multiple species, as shown in Figures 3 and 4. Generally, the flatmaps provide a simple 2D graphical representation for anatomical structures of a given species with overlays of nerve connections which enable the user to start exploring the entire organism. Flatmaps also provide details, where needed, to present an anatomically accurate picture of the connection between various parts of the body and the autonomic nervous system. Individual flatmaps are developed for different species, including human, rat, mouse, cat, and pig, providing (future) cross-species comparisons. Each flatmap consists of two main regions: the body including the visceral organs, and the central nervous system with more detail of the brain and spinal cord. These two regions are connected through directional edges to demonstrate neural connectivity associated with different biological systems and organs. The nerve fibres are categorized and rendered in the flatmap. The flatmaps also link between a given organ and anatomical structures with existing datasets at different biological scales through embedding of all existing experimental data and computational 3D scaffolds in relevant anatomical locations as discussed further in Section 7.

**Figure 4.**
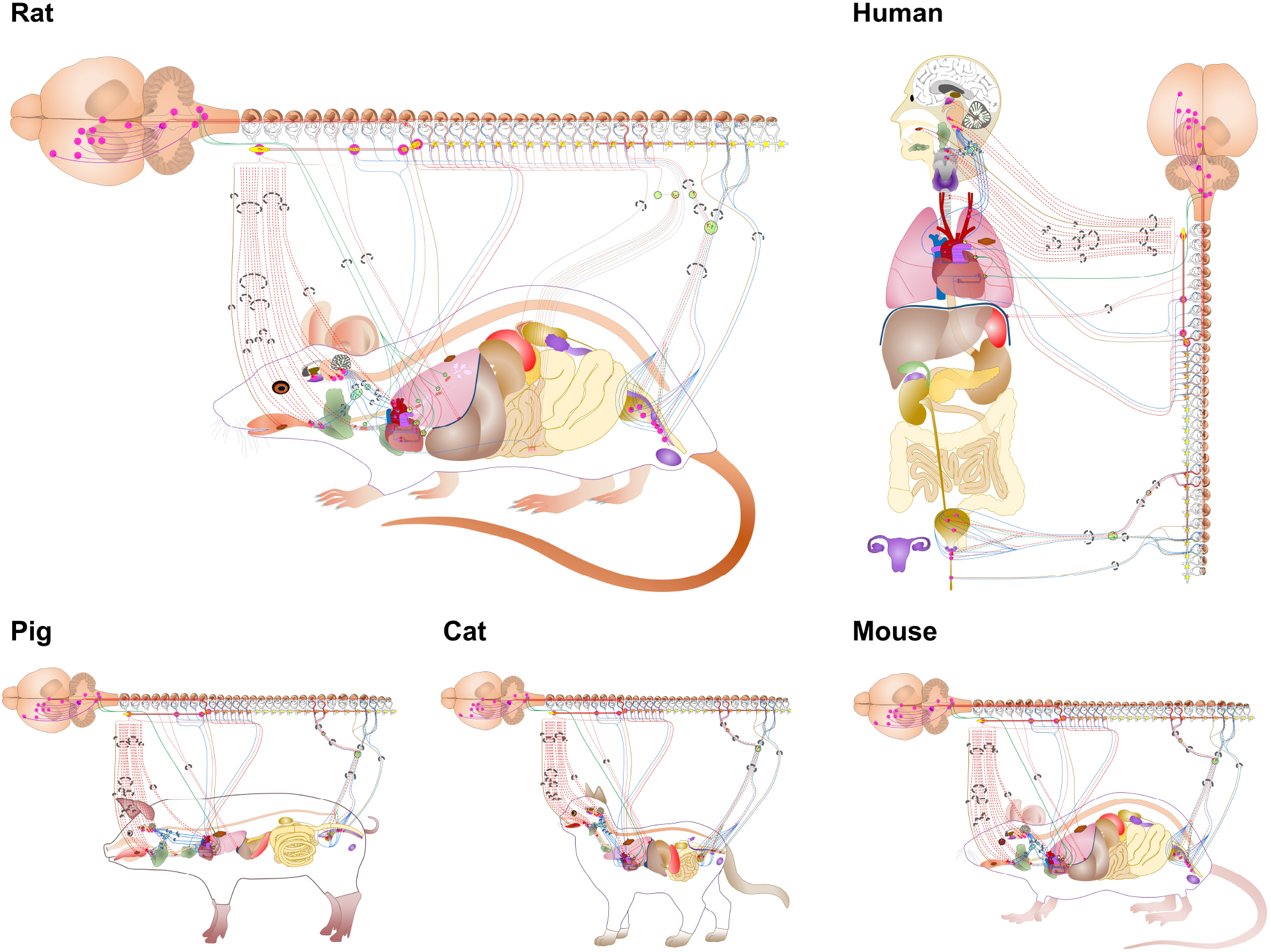
Flatmaps of individual species available through the SPARC portal at sparc.science/maps. Zooming in on these mapping diagrams provides additional levels of detail and clickable links that retrieve relevant SPARC data resources from Discover via the SKG (SPARC Knowledge Graph).

## 6 IMAGE SEGMENTATION AND ANNOTATION

We turn next to the processing of image data that are subsequently registered to the 3D anatomical scaffolds described in the following section. Neuroanatomical mapping experiments using image segmentation serve to inform scaffold models about the composition and trajectory of nerves and nerve-organ interactions. The diverse experimental paradigms for these mapping experiments span meso-to nano-resolution scales and use techniques such as microCT, tissue clearing, whole organ serial blockface image volumes, “flat-mount” organ preparations and electron microscopy. With each modality, a variety of segmentation goals, image formats and data sizes need to be accommodated. A FAIR image segmentation pipeline has therefore been established to enable the SPARC research program to harmonize experimental mapping data across different research groups. The keystones of this harmonization are a standard segmentation file format and rich metadata, collected and recorded within various data forms so that files can be understood on their own, as well as within the context of the dataset. In neuroanatomical mapping projects, file level metadata is targeted for capture for both segmentation files and the microscopy image file they are derived from.

The process of image segmentation creates digital models of the cellular and anatomical features of interest at the level of detail relevant for the research question at hand. Segmented anatomical elements are written in MBF Bioscience’s digital reconstruction file format. The format, in strides with SPARC’s FAIR and open data objectives, is documented in the Neuromorphological File Specification and is publicly available^24^ (Sullivan et al., 2020). A uniform data file format simplifies the registration procedure to a common coordinate space. Segmentations performed in other commercial, open-source, or custom software are converted to this common format during the curation process for scaffold fitting (Section 7). Generating mechanisms to speed data collection and segmentation file curation while reducing redundant data streams is a primary goal of the FAIR mapping effort.

Another key component of the SPARC program is the recognition of the importance of file-level information. This metadata is being carefully captured at the structure level for segmentations as well as within image data. Image segmentation software created by MBF Bioscience, e.g., Neurolucida 360^25^, has been specifically augmented to increase the quality and quantity of metadata inclusion while expanding the contributor base of datasets coming through the mapping pipeline. Access to species and organ-specific terminology within the software via connection to the SPARC ontology and vocabulary services within SciCrunch allows primary research annotators to select curated vocabulary terms and map neural, vascular and anatomical regions of interest. Utility of this interface within the segmentation software is dynamic and robust. It allows annotators to name cellular and anatomical features with community-accepted FAIR terminology while simultaneously enriching segmentation metadata with unique data identifiers specific to the term. For example, segmentation of the ventral region of the narrow open end of the mouse bladder could be identified as the “ventral bladder neck” – a term catalogued in the SKG and made available in MBF software. Annotation of this structure would then receive an ontological persistent identifier associated with the EMAPA ID space, part of the Edinburgh Mouse Atlas Project ontology^26^. This numerical identifier is stored in the segmentation file with the term “ventral bladder neck”, as well as the coordinate points that comprise the annotation. This structure-level metadata, in conjunction with subject-level metadata, allows for enhanced interoperability of experimental data on scaffold and flatmap models, and beyond. The tools that enable this style of data enrichment are available within MBF Bioscience segmentation software, regardless of the user’s involvement in the SPARC program, to encourage community-driven FAIR data production.

Bridges are provided to researchers from within the software to enable anatomy term requests for inclusion in the database accessed by the software. Term request pipelines have been established for SPARC researchers to strengthen vocabularies for organs, ganglia, nerve components, vasculature, and tissue. Entry points to that pipeline appear within MBF Bioscience software and directly on the SciCrunch SPARC Anatomy Working Group online dashboard. Once reviewed by the Anatomy Working Group and added to the database, the term is made available to the researcher the next time they use the software. The image segmentation workflow accounts for the dynamic changes to vocabularies within the SPARC ontology and vocabulary services within SciCrunch as the segmentation data files can be returned to at any point for review.

Cataloging relevant fiducial information (e.g., nerve roots, unique anatomical landmarks) using SPARC anatomical terminology is performed at multiple experimental stages in order to enable registration to the scaffolds and flatmaps. In conjunction with MAP-Core software engineers, MBF Bioscience engineers augmented the MBF Bioscience software interface to display static scaffold files. The static scaffold is created to replicate the experimental strategy used for segmentation (e.g. for flat-mount preparations, the scaffold is opened along the same cut lines as the experimental data). Researchers can display image data in conjunction with the representative static scaffold and are able to identify concordant data file fiducial points relative to the static scaffold. Once material coordinates for the segmentations are registered, the scaffold can be fitted to dynamic deformation models (described in the next section).

To enable further exploration and repurposing of image segmentation files from SPARC experimental datasets, we have developed an openly available application for file conversion of primary microscopy imaging data. The converter ingests proprietary image file formats and converts the files to more widely supported image formats, OME-TIFF and JPEG2000. The application enables the review and addition of consistent and rich metadata to increase the FAIRness of microscopy data in alignment with the metadata standards being developed by the SPARC consortium. The derived FAIR microscopy image and segmentation data, inclusive of rich experimental metadata, is curated by the SPARC Curation Team and supplied with the dataset on the SPARC Portal. A selection of derivative imaging data is selected by the researcher and prepared for viewing on the SPARC data portal, as MBF’s cloud-based image storing and viewing technology, Biolucida, is used to display image data on the Portal alongside the published dataset.

## 7 COMMON COORDINATE SYSTEMS AND 3D SCAFFOLDS

Many visceral organs of the body, including all of the high priority organs (heart, lungs, bladder, stomach, colon) for SPARC, undergo large deformations – hearts beat; lungs breathe; the bladder fills and empties; the stomach and colon, like the rest of the gut, are subject to large propagating waves of contraction. Even the kidneys and liver undergo small deformations associated with pulsatile blood flow. It is therefore imperative that any attempt to map the neurons within these organs must define the neurons and their cell bodies with respect to a three-dimensional (3D) material coordinate system within each organ (see Figure 5). The organs themselves also move with respect to the body that contains them, and the body moves with respect to the outside world. The embedding of organs within the body is described at the end of this section.

**Figure 5.**
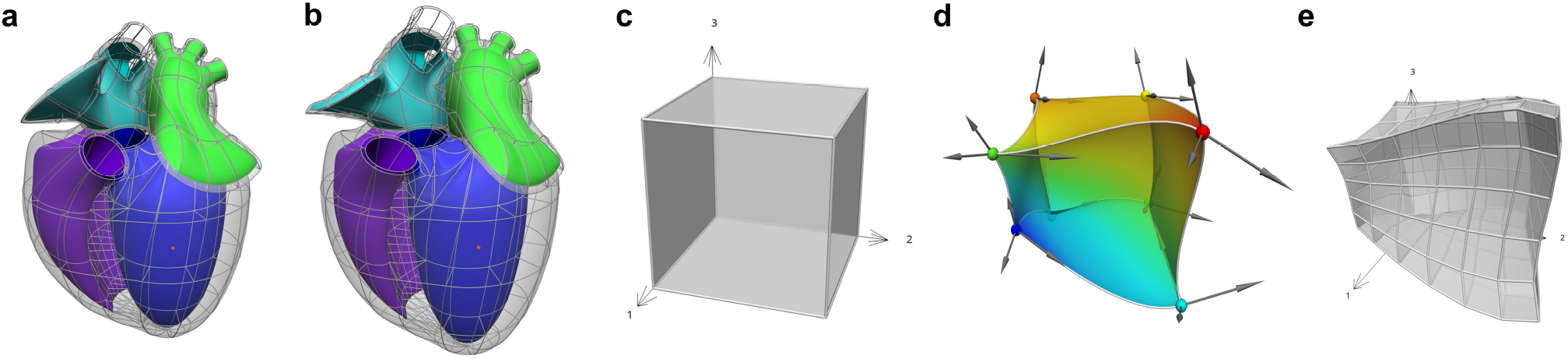
An Example of mapping from material space to physical space. (a-b) The heart scaffold defined with 282 tri-cubic Hermite elements. (c) A single element defined with (ξ_1_, ξ_2_, ξ_3_) coordinates. (d) The element in (c) mapped into 3D physical space where the colours illustrate a continuous scalar field defined over that space. In addition, (e) an example of a domain refinement of the deformed cube into subdomains or ‘elements’ is illustrated.

Anatomy is geometrically complex, and a very flexible approach is therefore needed to capture the intricacies of anatomical domains such as the heart, and lungs, etc. The Finite Element Method (FEM) that is widely used in the engineering world is a very powerful way to do this (Hunter and Smaill, 1988). The basic idea is that a physical domain (in this case an anatomical region of the body) is divided into subdomains (“elements”) joined via common nodes as illustrated in Figure 5 (this is called a finite element ‘mesh’). The field within an element is an interpolation using element basis functions of values of the field defined at the nodes (called the ‘nodal parameters’). The material space (defined by the element coordinates) can be refined to any degree in order to ensure that sufficient nodal parameters are available to match the resolution of the experimental data being mapped on that mesh.

An important feature of a 3D material coordinate system for organs is that it can deal with anatomical differences both within a species (individual differences) and across multiple species in order to enable cross-species comparisons. This can prove difficult when different species exhibit topological differences – as, for example, the varying numbers of pulmonary veins that enter the left atrium (typically 3 for rats, 4 for humans, and 2 for pigs). Another feature of mammals is that anatomical structures are generally smooth – there are none of the sharp edges typical of engineering structures.

To meet these requirements, we have developed Computer Aided Design (CAD) software called *Scaffold-Maker*^27^ specifically for creating 3D material coordinate systems for body organs. Note that the term ‘material’ is used because these coordinates effectively identify the position of any material (tissue) particle, independent of its location in 3D space or how distorted it is. We could use the term ‘tissue coordinates’ rather than ‘material coordinates’ but the idea is more general – for example, a material coordinate position inside a deforming human body locates a unique material point in the body but this is not necessarily part of a tissue (e.g. it could be in the middle of an airway). We call the 3D material coordinate system for an organ, a *‘scaffold’*, because it is a coordinate framework into which many different aspects of tissue structure can be assembled, including muscle fibre orientations, vascular geometry, neural pathways and the spatial distributions of RNA-Seq data. The anatomical scaffolds are generated from connected, simple-shaped elements to follow the features of an organ or other anatomical part. A combination of element plus local coordinates within the scaffold gives a material coordinate system labelling the same piece of tissue across all configurations for an individual, or an equivalent anatomical location across multiple individuals.

Each scaffold is created with an idealized reference geometry close to the expected *in-vivo* configuration of the organ. It defines distinct regions and landmarks annotated with standard anatomical terms enabling their correspondence with data digitised from specimens in experiments.

Here it is worth emphasizing here the concept of a *field,* which is a function that maps all the material coordinates of the scaffold (its domain) to a set of values. The scaffold’s geometry is defined by a coordinate field mapping into an *x,y,z* Cartesian coordinate system, which is special in that it maps each material point (which can have multiple labels on element boundaries) to distinct locations in the coordinate system consistent with the connectivity and continuity within and between its elements. Figure 5 illustrates the idea of common material coordinates in a physical space using two different configurations of a heart scaffold. A material point is always identified by the same material coordinates even when the tissue is moved and stretched. Any functions can be used to define a field, however we use *C*^1^-continuous cubic Hermite polynomial *basis functions* to map the geometry which is the simplest function able to give first derivative continuity giving the smooth shapes exhibited by most organs and parts of the body (Bradley et al., 1997; Fernandez et al., 2004).

Once these mappings are established, a scalar or vector field can be represented on the material spaces of Figure 5a-b, using basis functions that span the normalised xi¿-coordinates, and are mapped to the physical domains of Figure 5a-b, as illustrated for a scalar field on a 3D domain by the spatially varying colours in Figure 5d.

We illustrate the application of these concepts to building scaffolds for the human, pig, and mouse colon in Figure 6. First, a sub-scaffold (Figure 6a-c) is built that captures the anatomy of the haustra (the bulging wall segments) and taeniae coli (the longitudinal muscle bands) for each species – note the species differences (the human has 3 taeniae coli, the pig two and the mouse none). Next, the colon centerline is defined (Figure 6d-f) – again note the very different centerlines for the three different species. Finally, the sub-scaffolds are algorithmically attached sequentially along the centerline to form the final scaffold for each species (Figure 6g-i).

**Figure 6.**
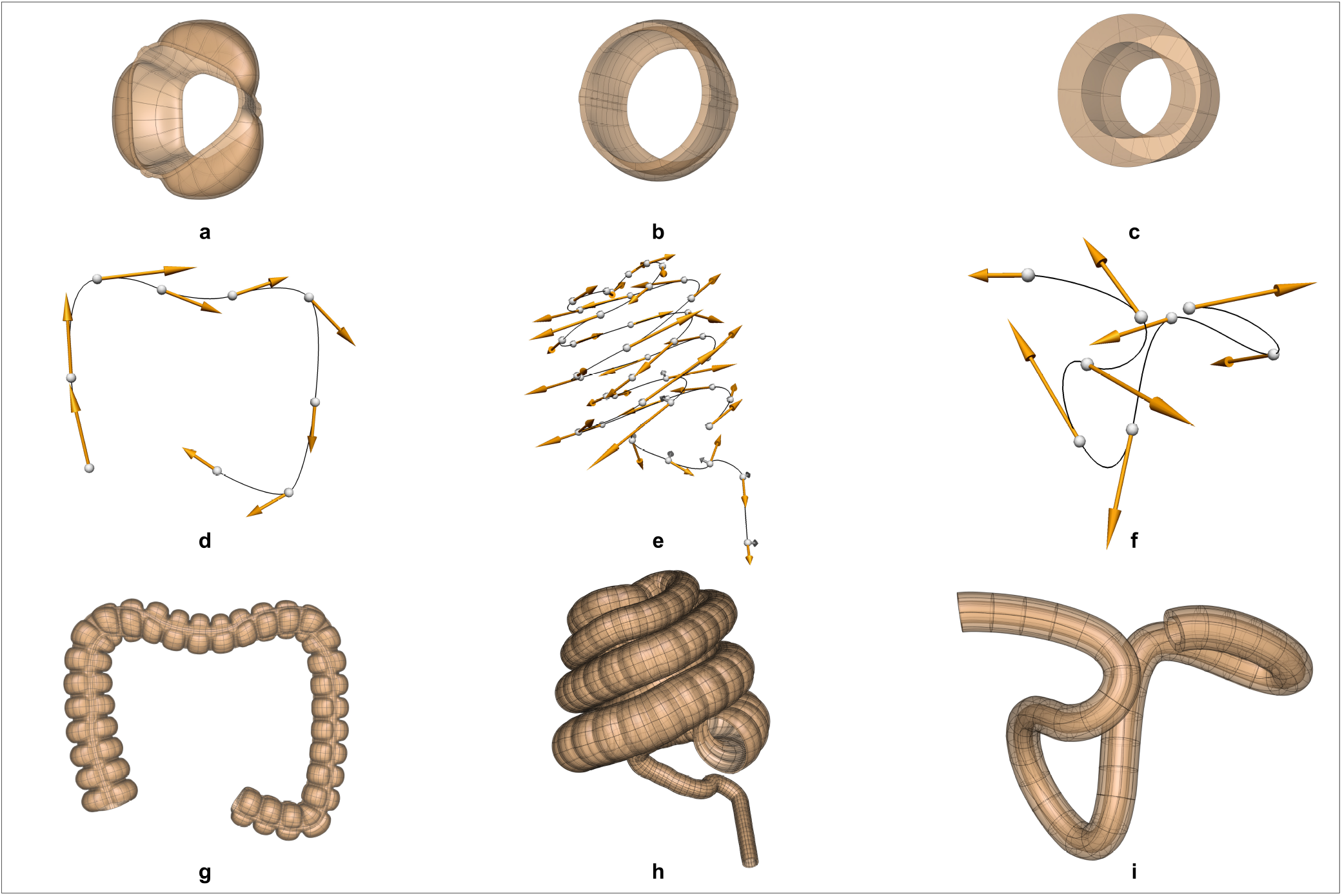
Scaffolds for the colon in the human, pig and mouse. The scaffolds for each species are built in three steps: (a-c) define a sub-scaffold that captures the cross-sectional anatomy of the colon for the three species, (d-f) define the centerline of the colon, (g-i) attach these sub-scaffolds sequentially to the centerline to form the final scaffold.

For each experimental specimen, a *fitting* process computes coordinate field parameters for the scaffold which give an optimal correspondence to the annotated coordinate data digitized from the specimen. Proper fitting must closely conform to landmarks and boundaries of anatomical features, but otherwise smoothly spread out the scaffold geometry in between fixed landmarks in proportion to the reference coordinate state. While it is approximate, this process gives confidence that the same material coordinates on the scaffold refer to equivalent anatomical locations in all specimens across the study.

In some cases, the topology of the scaffold depends on the species. For example, heart scaffolds need four pulmonary veins entering the left atrium in the human heart (Figure 7a), two for the pig heart (Figure 7b) and three for the rat heart (Figure 7c).

**Figure 7.**
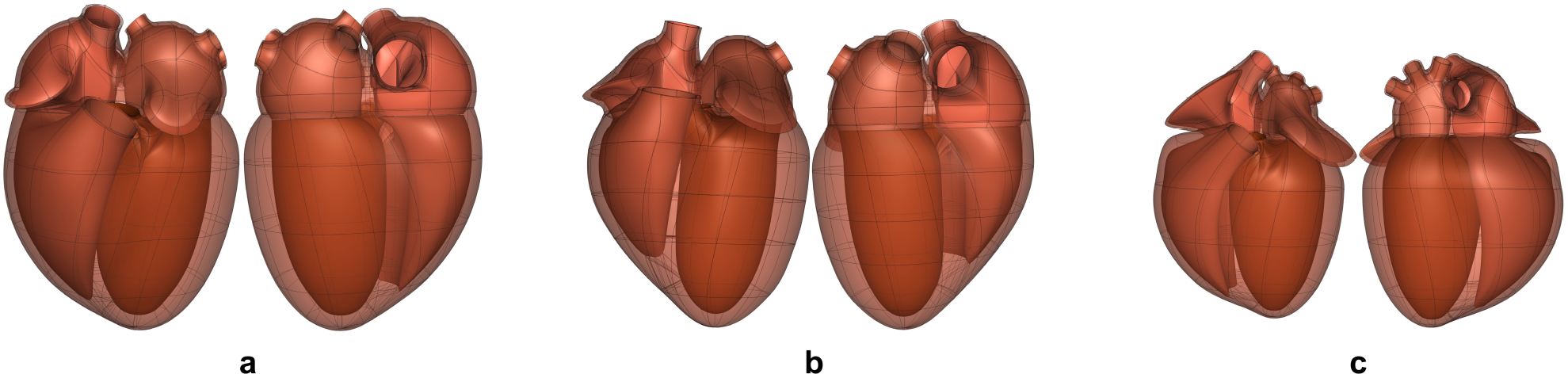
Front (left) and back (right) views of the (a) human, (b) pig and (c) rat scaffolds (not to scale). The topology of the scaffold is different for each species as each has a different number of pulmonary veins entering the left atrium (4, 2, and 3, respectively).

With the scaffold in the fitted geometric configuration for a specimen, all other data such as digitized neurons and pathways in that configuration can be in material coordinates, or data can be used to fit continuous fields. Once this data registration is complete, data from multiple specimens can be compared in any common coordinate system over the scaffold, for example the idealized reference coordinates for the scaffold (Leung et al., 2020), or the average coordinates across multiple specimens (Osanlouy et al., 2020).

### 7.1 Integration with whole body models

The organ scaffolds and neural pathways can also be located within the 3D body coordinates of the intact animal, as illustrated for the rat in Figure 8. The 3D body coordinate system defines material coordinates that can map to multiple species and to multiple instances of a particular species in order to facilitate comparisons across species or across instances within a species. We have defined a cylindrical core that contains the visceral organs and a layer that includes the spinal cord and can be extended to define the limbs. The cylindrical core is divided by the diaphragm into the thorax and abdomen. The core and outer layer can be subdivided into an arbitrary number of segments in order to capture the desired level of anatomical accuracy. An organ scaffold is located within the body scaffold by defining a small number of fiducial points for each organ that ensure that when the scaffold is fitted to body surface data, it also matches key anatomical points (e.g. the diaphragm, the aortic root, etc) to ensure that they are positioned correctly relative to other parts of the body. The data fitting for this simple example uses the outer skin surface, the diaphragm, and the centerline of the spinal cord.

**Figure 8.**
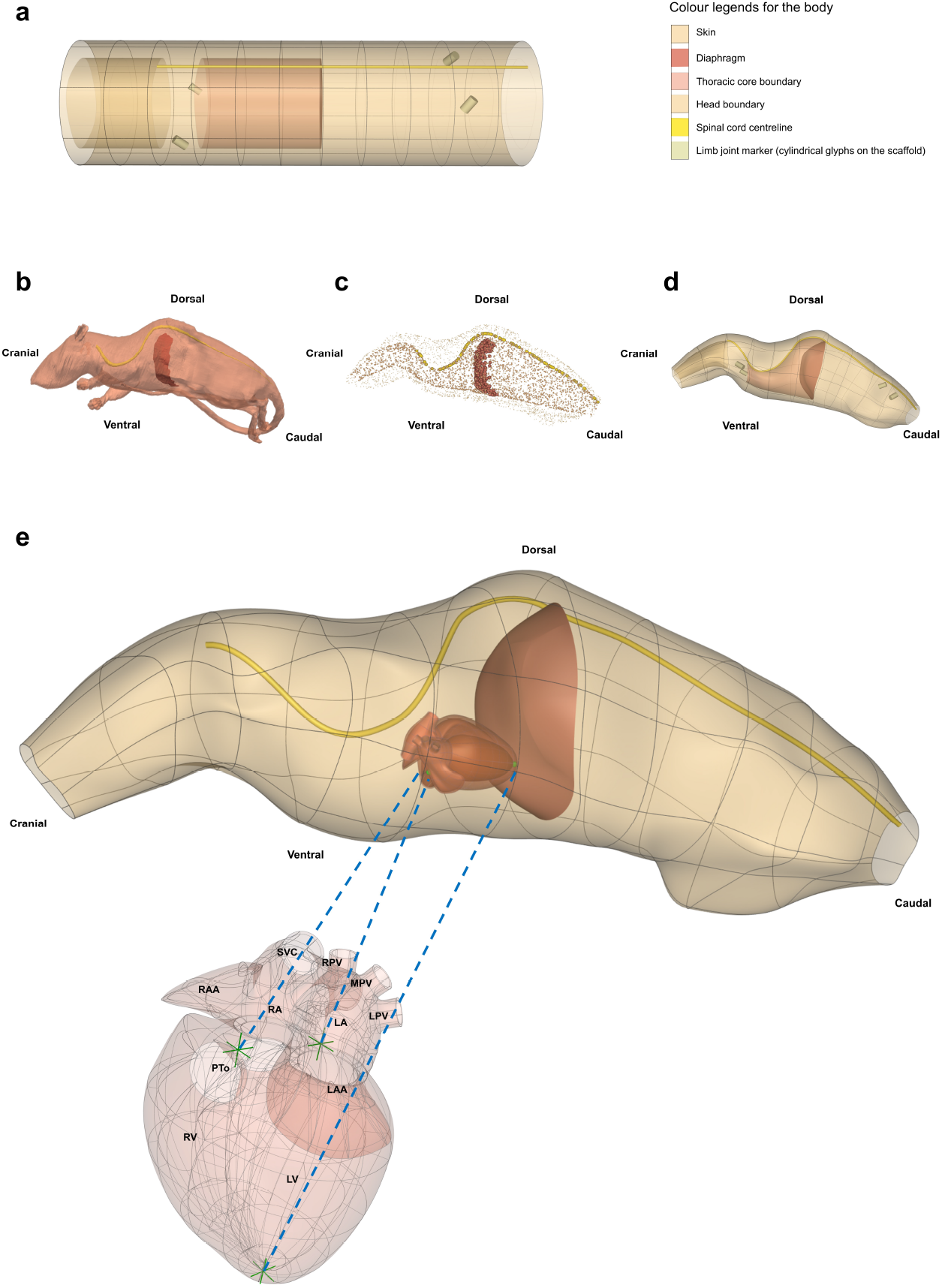
Generating a whole rat body anatomical scaffold from data for integration with other organ scaffolds. (a) The 3D *body coordinate system* shows distinct anatomical regions with different colours. (b) The rat model (NeuroRat, IT’IS Foundation. DOI: 10.13099/VIP91106-04-0) in the vtk format with spinal cord and diaphragm visible. This model contains 179 segmented tissues with neuro-functionalized nerve trajectories (i.e., associated electrophysiological fiber models). (c) All tissue tissues from the rat body were resampled and converted into a data cloud to provide the means for generating the whole body scaffold. The data cloud is a 3D spatial representation of the tissue surface consisting of a certain density of points in the 3D Euclidean space. These points are used to define the objective function for the scaffold fitting procedure. The data cloud of the skin, inner core, spinal cord, and diaphragm are shown in (c) with the limbs and tail excluded. (d) The 3D body scaffold (a) was fitted to the rat data to generate the anatomical scaffold. (e) A generic rat heart scaffold was projected into the body scaffold using three corresponding fiducial landmarks (green arrows in heart and green spheres in the body). This transformation allows embedding of the organ scaffolds into their appropriate locations in the body scaffold, providing the required physical environment required for simulation and modeling. SCV: superior vena cava; RPV: right pulmonary vein; MPV: middle pulmonary vein; LPV: left pulmonary vein; RAA: right atrial appendage; RA: right atrium; LA: left atrium; LAA: left atrial appendage; PTo: pulmonary trunk outlet; RV: right ventricle; LV: left ventricle.

When computational models feature anatomy representations or have corresponding anatomical locations, they are integrated within whole-body anatomical models (NEUROCOUPLE and NEUROFAUNA) and/or 3D organ scaffolds. Integrating computational models within multi-scale anatomical models (‘model functionalization’) achieves a range of purposes (Neufeld et al., 2018). It provides the physical environment required to simulate the interaction between devices and the organism (e.g., a dielectric environment that is exposed to an electroceutical implant, or an acoustic environment in which neuromodulation is performed with ultrasound stimulation), it facilitates the identification of colocalized data and models (e.g., to find data suitable as model input, or for model validation, or to identify two models that might be coupled), and it provides a natural, anatomy-driven means of searching for data and models. For whole-body model-integration and to capture the anatomical representation of ANS connectivity (complementing the functional and relational representation in ApiNATOMY and flatmaps), detailed whole-body anatomical models have been created through image-segmentation – with particular attention to the ANS – and functionalized with electrophysiological nerve fiber models. The NEUROCOUPLE human male and female models are based on cryosection images – color photographs of frozen sectioned cadavers – of unique resolution and contrast (Park et al., 2006). Around 1200 different anatomical regions, including in excess of 200 nerves and 900 nerve trajectories, more than 320 muscles, as well as many vascular segments are distinguished. The NEUROFAUNA rat model was segmented from multiple MRI and MicroCT scan sequences with optimized contrast variations, in order to visualize and accurately delineate important tissue types and organs, as well as bones. The whole-body rat model has been functionalized with nerve trajectories, and the detailed Waxholm brain atlas co-registered to the image data (Papp et al., 2014).

## 8 EXAMPLE OF DATA CURATION, SEGMENTATION AND REGISTRATION

We illustrate the SPARC DRC data workflow by showing how measurements of intrinsic cardiac neurons (ICNs) from the DISCOVER data repository^28^ are mapped into a heart atrial scaffold (Figure 9). The ICNs are a local, integrative neural circuitry system which mediates and regulates the vagus nerve’s control of the heart. Accurate characterization of these neurons requires a comprehensive quantification of their spatial distribution on the heart. Data was acquired using an experimental workflow that uses a Knife Edge Scanning Microscope (KESM) technology to acquire high-resolution images of the rat heart tissue. Cardiac regions and single ICN were segmented and annotated using the MBF Tissue Mapper software^29^. These data were used to generate and customize individual heart scaffolds to enable the registration of ICNs (Leung et al., 2020). Since ICNs are mainly distributed around the basal part of the heart and pulmonary veins, a ‘generic’ scaffold was algorithmically generated from a set of anatomical and mathematical parameters for the atrial topology using the Scaffold-Maker software. This atrial scaffold consists of separate manifolds to represent different anatomical regions enclosing both the endocardial and epicardial tissue volumes: left atrium (LA), left atrial auricle (LAA), left pulmonary vein (LPV), middle pulmonary vein (MPV), right pulmonary vein (RPV), right atrium (RA), right atrial auricle (RAA), inferior vena cava (IVC), and superior vena cava (SVC). The separation of these structures into distinct groups were necessary for an effective fitting of the scaffold to data and accurate mapping of the ICNs.

**Figure 9.**
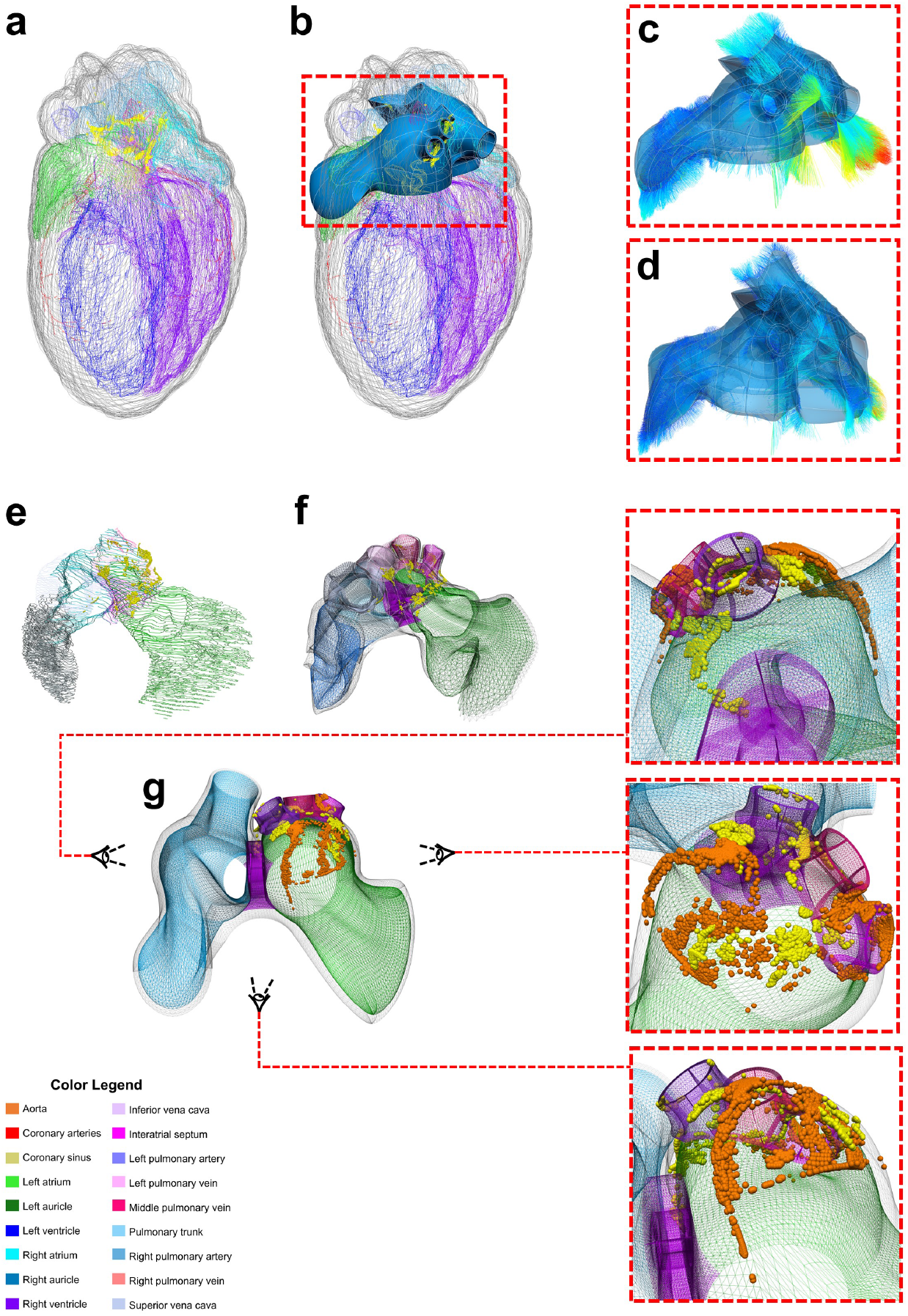
Mapping individual rat ICNs and cardiac anatomy onto a generalized 3D scaffold for comparison across animals through an automated pipeline. (a) Segmentation contours from the imaged heart is used to (b) customise and generate the atrial scaffold; (c) individual data points from the contours are projected on the nearest scaffold surface to compute the minimum distance for the fitting process to (d) fit and morph the scaffold to match the data; (e) the atrial contour data with ICN cells are processed to (f) map and register the spatial distribution of individual cells on the scaffold as material points. This material point mapping provides a powerful way to (g) capture information from multiple subjects (e.g. orange vs. yellow spheres showing individual ICNs from two different rats) on one ‘generic’ scaffold. Here the heart is visualized from a superior angle to appreciate how the neuron locations in the original data are projected onto the generic scaffold. Figure adapted from (Leung et al., 2020).

Experimental preparations of the heart tissue for imaging often lead to distortions and deformations of the organ and thus may not be reflective of true heart geometry *in situ.* Fitting a scaffold to this data allows the preservation of the local material coordinates of the heart tissue, which can then be used to recover the generic geometry of the heart. Therefore, the customized generic atrial scaffold was fitted to individual subject’s image segmentation contours using a linear least squares optimization. Specifically, the sum of the weighted distances between each segmentation data point on the contour and its projection onto the nearest element was minimized during the fitting process. This distance is a function of the scaffold nodal parameters. The projected points are computed from the nodal parameter interpolation; the global coordinates of the projected points are a function of local material coordinates i.e. ξ_1_, ξ_2_, and ξ_3_ and their partial derivatives. To avoid scaffold shape distortion in this process, a term was introduced to the optimization to penalise for the cardiac tissue strain. Additionally, a smoothness constraint was often added in the fitting process to measure the deformation of the surface and regularize the problem since segmentation contours were sometimes either insufficient or noisy in different regions.

The fitted scaffold can capture the spatial distribution of ICN cells and embed them locally into the elements. The material embedding of ICNs stores a unique one-to-one mapping that can be used to transform the cells onto the corresponding generic scaffold elements. The generic scaffold offers the opportunity to register ICNs from different rat samples onto one ‘integrative’ scaffold. Integration of multiple datasets allows for quantitative comparison across multiple species despite variation in the coordinate system and topographic organization of neurons.

## 9 COMPUTATIONAL MODELING AND DATA ANALYSIS

An online-accessible open source computational modeling platform called o^2^S^2^PARC (‘open, online simulations for SPARC’; Fig. 10) has been developed and is available through a web interface under the *MIT license.* The principal considerations were to enable shared modeling and data analysis activities in order to make software modules developed by individual research groups as reusable and interoperable as possible. Curated and published computational studies are available on the SPARC portal.

**Figure 10.**
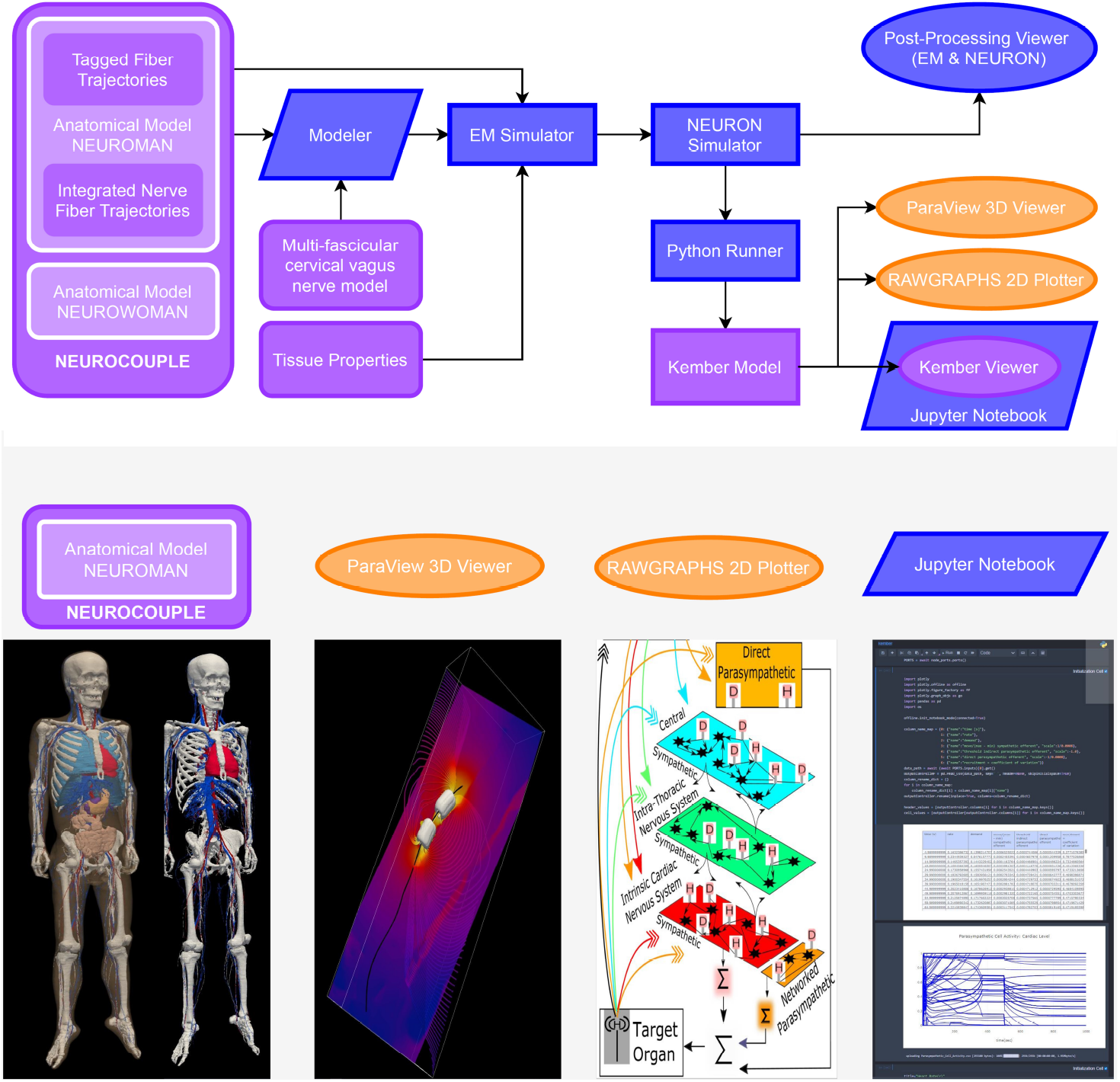
Biophysical modeling and analysis workflow implemented at the end of the first o^2^S^2^PARC development year as a demonstrator of the feasibility of an open, readily extensible, online-based platform for collaborative computational neurosciences. (purple: published on the portal, blue: o^2^S^2^PARC services, yellow: 3rd party viewers embedded in o^2^S^2^PARC) Electromagnetic exposure of the vagus nerve by a bioelectronic implant is simulated, along with the resulting vagus nerve stimulation and its impact on cardiovascular activity: A ‘NEUROCOUPLE’ service injects the male NEURONMAN anatomical model, along with a multi-fascicular cervical vagus nerve model layer and integrated nerve fiber trajectories, into a ‘Modeler’ service, where interactive constructive geometry is used to integrate an implant electrode geometry. The different NEUROMAN anatomical regions are already tagged with tissue names and the fiber trajectories with electrophysiological information (fiber type and diameter). The implant-enhanced anatomical model serves as input to an ‘EM Simulator’ service, which obtains dielectric properties from a ‘Tissue Properties’ service and assigns them to regions based on their tags. After defining boundary conditions, discretization, and solver parameters, the high-performance computing enabled electro-quasistatic solver is called. The resulting electric potential is injected into a ‘NEURON Simulator’ service, which also receives the tagged fiber trajectories from the ‘NEUROCOUPLE’ service as input and discretizes these trajectories into compartment with associated ion channel distributions (according to predefined, diameter-parameterized fiber models), produces input files for the neuronal dynamics simulator from NEURON, and executes the simulations using NEURON’s ‘extracellular potential’ mechanism to consider the electric field exposure. A ‘Python Runner’ service implements the computation of stimulation selectivity indices, which feed a ‘Kember Model’ service that contains an implementation of the multi-scale cardiac regulation model from (Kember et al., 2017). A ‘Paraview 3D Viewer’ service and a ‘Rawgraphs Viewer’ service provide result visualization. Finally, a ‘Kember Viewer’ service performs predefined post-processing analysis on the output of the ‘Kember Model’ service and visualizes the results. The ‘Kember Viewer’ service is a specialized instance of the ‘Jupyter Notebook’ service, which permits to present a (Python) script along with documentation in an interactively explorable and editable form.

The platform is deployed on dedicated SPARC servers, as well as scalably in the cloud via Amazon Web Services (AWS) to provide access to computational resources such as GPU clusters, which might not be readily available to some users. Users access it with an associated Python client, or can use a *REST* API, which follows the *OpenAPI* standard. It can also be deployed on a local machine or cluster. The online platform provides the user with an overview of available data and computational resources (SPARC data on Blackfynn *DISCOVER,* as well as S3 storage repositories), published computational services to which the user has access, and to owned and shared computational studies (projects) and study templates (pre-setup study-structures that represent common workflows that can be instantiated and modified as needed). When creating a new study or accessing an existing one, a workbench view illustrates the graph (pipeline) of connected nodes (computational or interactive services) – outputs of nodes can be passed as input to other nodes and connected to specific ports. The hierarchical structure permits nodes to contain nested sub-graphs. Sub-graphs can be packaged as macro’s (with fine-grained control over the exposed ports) and shared with the community as reusable building-blocks. Nodes can be accessed, upon which a common GUI for specifying parameters or connecting available inputs from upstream nodes to ports is displayed (this GUI is automatically generated based on the provided meta-data – see below), along with a flexibly configurable *iframe* that can be used for visualization and interaction, and over which the service can have full control. A guided mode permits to wrap expert-developed workflows into a wizard-like step-by-step application that hides the underlying complexity and only exposes parameters and interactions that require user input.

Services are packaged as Docker images around which o^2^S^2^PARC provides an extensive framework. Docker technology offers lightweight, rapidly deployable environments (similar to *virtual machines*). While the user is responsible for providing the image content, the framework automatically takes care of providing implementation-agnostic interoperability and seamless integration into the platform, including logging, interservice communication and communication with the web front-end, data collection and forwarding, as well as job scheduling and execution on the appropriate hardware respecting GPU, CPU and RAM requirements. Using Docker images offers large flexibility for the content (operating system, software environments, etc.) and permits packaging of software along with the environment under which it was developed and tested (for sustainability and reproducibility reasons; users do not have to worry about compatibility, software dependencies, and installation requirements). Reproducibility is further advanced by versioning changes in evolving services, such that a user can choose to re-execute a study using the original service versions, the newest available versions, or a combination of manually selected versions. Examples of published services available in o^2^S^2^PARC include simulators (e.g., *in vivo* electromagnetic field computation, neuronal dynamics), organ and tissue electrophysiology models (e.g., multiscale models of cardiac neuro-regulation), data and model viewers, machine-learning and data analysis modules (e.g., histology segmentation, spike sorting), etc. A particularly interesting service class are script-based services, such as ‘runners’, which combine executable scripts (Python, Octave, CellML, etc.) with suitable interpreters as black-box modules, or interactive environments that present them as viewable/editable and executable code (e.g., JupyterLab). Such services are valuable for the publication, reuse, sharing, and joint elaboration of scripts, e.g., for data analysis. For an example study, see Figure 10.

Quality assurance in the development of o^2^S^2^PARC is ascertained through compulsory code-reviews, automatic code analysis and dependency updates (e.g. *codeclimate* and *dependabot*), unit-, regression-, and end-to-end-testing (e.g., with *puppeteer*), multi-staged release (development, staging, and production versions), while service QA is established providing a template mechanism (*cookiecutter*) that includes automatic testing of communication interfaces, support for the co-submission of test benchmarks. Supporting information on verification and validation can be attached to services and studies as part of displayed meta-data (complementing the compulsory meta-data from the cMIS), which is part of the existing and forthcoming functionality to facilitate, encourage, and enforce adoption of the ‘Ten Simple Rules’ (Erdemir et al., 2020).

## 10 DISCUSSION AND CONCLUSIONS

The SPARC program is designed to deliver a comprehensive and quantitative description of the mammalian autonomic nervous system via a web portal that provides access to experimental data, neural connectivity maps, anatomical and functional models, and computational resources. The role of the four cores of the SPARC DRC is to curate, annotate and store data and models, to enable sharing of reproducible analyses, to map data from multiple species into common coordinate anatomical frameworks so that they can be compared and made relevant to human physiology, and to enhance our understanding of the autonomic nervous system via computational models that simulate its physiological function and can be used to devise and optimize new bioelectronic treatment approaches. The success of the project will be measured by the extent to which these data, models and tools contribute both to our scientific understanding of autonomic function and to the design of neuromodulation devices that improve healthcare outcomes by targeting autonomic dysfunction.

A critical aspect of data management for the SPARC program has been the development and implementation of minimum information standards and curation/annotation workflows that ensure metadata is available to support FAIR data principles. This is a challenging task, both for the DRC and for the SPARC experimental groups that are having to adapt to new best practice, but delivering reproducible and reusable data, data analyses, and computational models is an important and essential outcome of the project with significance well beyond autonomic neuroscience. Similarly, the concepts implemented as part of the o^2^S^2^PARC platform to address the quality assurance, reproducibility, compatibility and sustainability challenges mentioned in Section 2 are expected to be readily applicable and valuable to the wider computational modeling and analysis community.

As well as the continued curation, annotation, and mapping of datasets provided by the SPARC experimental community, future developments of the SPARC DRC infrastructure include: (i) more complete neural connectivity displayed on the flatmaps and more automated ways of drawing these pathways directly from the semantic annotations deposited into the SPARC Knowledge Graph hosted within SciCrunch; (ii) further organ scaffolds beyond the current set (heart, lungs, stomach, bladder, colon); (iii) whole body scaffolds for multiple species, into which the organ scaffolds and neural pathways can be algorithmically embedded; (iv) further development of tools to enable experimentalists to register their data into the organ and body scaffolds; (v) implementation of built-in quality assurance functionality; (vi) provision of extended data processing functionalities and support for sharing and publishing analyses; (vii) facilitation of *in silico* studies and provision of expert workflows; and (viii) an increasing focus on the interpretation of autonomic nervous system connectivity with models that explain function in order to facilitate the design, optimization, safety assessment, personalization, and control of effective device-based neuromodulation therapies.

## ACKNOWLEDGMENTS

The authors would like to gratefully acknowledge all of their SPARC colleagues for their contributions to the SPARC DRC program, including the SPARC experimental groups and the SPARC NIH program officers, all of whom have contributed to the work presented here.

## AUTHOR CONTRIBUTIONS

M. Osanlouy from MAP-Core leads the cardiac data registration work, generated a number of the figures and helped with overall coordination of the paper. B. de Bono from MAP-Core and K-Core leads the development of ApiNATOMY. D. Brooks and N. Ebrahimi from MAP-Core lead the flatmap visualisation. R. Christie from MAP-Core leads the 3D scaffold developments. M. Lin from MAP-Core leads the colon data registration. D. Nickerson is Technical Lead for MAP-Core. E. Soltani from MAP-Core leads the 3D body scaffold work. A. Bandrowski, J. Grethe, T. Gillespie, and M. Martone from MAP-Core have all participated in the development of the SPARC Data Structure and all have contributed to the writing and revision of this manuscript. S. Tappan from MAP-Core leads MBF Bioscience software development in support of SPARC program goals and has contributed to paper writing. M. Heal from MAP-Core has contributed to paper writing. J. Wagenaar is the principal investigator of the DAT-Core and leads the development of the data management platform. L Guercio is the project manager of the DAT-Core, has contributed to writing this paper and is actively involved in development of the platform and SPARC Portal. N. Kuster is the Principal Investigator leading the SIM-CORE. E. Neufeld from SIM-CORE leads the o^2^S^2^PARC platform design and development and has contributed to the paper writing. K. Zhuang from SIM-CORE has contributed to standardization, the described applications, and paper illustration. A. Cassara from SIM-CORE has contributed to the computational modeling. P. Hunter, who is the PI for MAP-Core led the overall coordination of writing for the paper.

## Footnotes

1 https://commonfund.nih.gov/sparc

2 https://sparc.science/

3 https://www.go-fair.org/fair-principles

4 https://discover.blackfynn.com

5 https://scicrunch.org

6 https://www.mbfbioscience.com/biolucida

7 https://osparc.io

8 https://discover.blackfynn.com

9 https://www.go-fair.org/fair-principles

10 https://github.com/bvhpatel/SODA/wiki

11 https://github.com/SciCrunch/NIF-Ontology/tree/sparc

12 https://www.protocols.io

13 https://github.com/bvhpatel/SODA/wiki

14 https://braininitiative.nih.gov/

15 https://www.mbfbioscience.com/microfileplus

16 https://neuinfo.org

17 https://scicrunch.org/scicrunch/interlex/dashboard

18 https://github.com/SciGraph/SciGraph

19 https://www.fdilab.org/scicrunch

20 http://physiomeproject.org/

21 https://models.physiomeproject.org/welcome

22 https://www.cellml.org

23 https://github.com/SciCrunch/NIF-Ontology/tree/sparc

24 www.mbfbioscience.com/filespecification

25 https://www.mbfbioscience.com/neurolucida360

26 https://www.emouseatlas.org/emap/home.html

27 https://github.com/ABI-Software/scaffoldmaker

28 https://discover.blackfynn.com/datasets/77

29 https://www.mbfbioscience.com/tissue-mapper

## REFERENCES

Bandrowski, A., Grethe, J. S., Pilko, A., Gillespie, T. H., Pine, G., Patel, B., et al. (2021). Sparc data structure: Rationale and design of a fair standard for biomedical research data. bioRxiv

Bradley, C., Pullan, A., and Hunter, P. (1997). Geometric modeling of the human torso using cubic hermite elements. Annals of biomedical engineering 25, 96–111

Bug, W. J., Ascoli, G. A., Grethe, J. S., Gupta, A., Fennema-Notestine, C., Laird, A. R., et al. (2008). The nifstd and birnlex vocabularies: building comprehensive ontologies for neuroscience. Neuroinformatics 6, 175–194

De Bono, B., Grenon, P., and Sammut, S. J. (2012). Apinatomy: A novel toolkit for visualizing multiscale anatomy schematics with phenotype-related information. Human mutation 33, 837–848

De Bono, B., Safaei, S., Grenon, P., and Hunter, P. (2018). Meeting the multiscale challenge: representing physiology processes over apinatomy circuits using bond graphs. Interface focus 8, 20170026

Erdemir, A., Mulugeta, L., Ku, J. P., Drach, A., Horner, M., Morrison, T. M., et al. (2020). Credible practice of modeling and simulation in healthcare: ten rules from a multidisciplinary perspective. Journal of translational medicine 18, 1–18

Fenner, M., Crosas, M., Grethe, J. S., Kennedy, D., Hermjakob, H., Rocca-Serra, P., et al. (2019). A data citation roadmap for scholarly data repositories. Scientific data 6, 1–9

Fernandez, J., Mithraratne, P., Thrupp, S., Tawhai, M., and Hunter, P. (2004). Anatomically based geometric modelling of the musculo-skeletal system and other organs. Biomechanics and modeling in mechanobiology 2, 139–155

Garny, A. and Hunter, P. J. (2015). Opencor: a modular and interoperable approach to computational biology. Frontiers in physiology 6, 26

Gorgolewski, K. J., Auer, T., Calhoun, V. D., Craddock, R. C., Das, S., Duff, E. P., et al. (2016). The brain imaging data structure, a format for organizing and describing outputs of neuroimaging experiments. Scientific data 3, 1–9

Hunter, P. J. and Borg, T. K. (2003). Integration from proteins to organs: the physiome project. Nature reviews Molecular cell biology 4, 237–243

Hunter, P. J. and Smaill, B. H. (1988). The analysis of cardiac function: a continuum approach. Progress in biophysics and molecular biology 52, 101–164

Kember, G., Ardell, J. L., Shivkumar, K., and Armour, J. A. (2017). Recurrent myocardial infarction: mechanisms of free-floating adaptation and autonomic derangement in networked cardiac neural control. PloS one 12, e0180194

Leung, C., Robbins, S., Moss, A., Heal, M., Osanlouy, M., Christie, R., et al. (2020). 3d single cell scale anatomical map of sex-dependent variability of the rat intrinsic cardiac nervous system. BioRxiv doi:10.1101/2020.07.29.227538

Lloyd, C. M., Halstead, M. D., and Nielsen, P. F. (2004). Cellml: its future, present and past. Progress in biophysics and molecular biology 85, 433–450

Low, P. A. (2011). Primer on the autonomic nervous system (Academic Press)

Mungall, C. J., Torniai, C., Gkoutos, G. V., Lewis, S. E., and Haendel, M. A. (2012). Uberon, an integrative multi-species anatomy ontology. Genome biology 13, 1–20

Neufeld, E., Lloyd, B., Schneider, B., Kainz, W., and Kuster, N. (2018). Functionalized anatomical models for computational life sciences. Frontiers in physiology 9, 1594

Osanlouy, M., Clark, A. R., Kumar, H., King, C., Wilsher, M. L., Milne, D. G., et al. (2020). Lung and fissure shape is associated with age in healthy never-smoking adults aged 20–90 years. Scientific Reports 10, 1–13

Papp, E. A., Leergaard, T. B., Calabrese, E., Johnson, G. A., and Bjaalie, J. G. (2014). Waxholm space atlas of the sprague dawley rat brain. Neuroimage 97, 374–386

Park, J. S., Chung, M. S., Hwang, S. B., Shin, B.-S., and Park, H. S. (2006). Visible korean human: its techniques and applications. Clinical Anatomy: The Official Journal of the American Association of Clinical Anatomists and the British Association of Clinical Anatomists 19, 216–224

Starr, J., Castro, E., Crosas, M., Dumontier, M., Downs, R. R., Duerr, R., et al. (2015). Achieving human and machine accessibility of cited data in scholarly publications. PeerJ Computer Science 1, e1

Sullivan, A. E., Tappan, S. J., Angstman, P. J., Rodriguez, A., Thomas, G. C., Hoppes, D. M., et al. (2020). A comprehensive, fair file format for neuroanatomical structure modeling. bioRxiv

Wilkinson, M. D., Dumontier, M., Aalbersberg, I. J., Appleton, G., Axton, M., Baak, A., et al. (2016). The fair guiding principles for scientific data management and stewardship. Scientific data 3, 1–9

